# Hybrid Population PK-Machine Learning Modeling to Predict Infliximab Pharmacokinetics in Pediatric and Young Adult Patients with Crohn’s Disease

**DOI:** 10.1101/2025.05.01.651780

**Authors:** Kei Irie, Phillip Minar, Jack Reifenberg, Brendan M Boyle, Joshua D Noe, Jeffrey S Hyams, Tomoyuki Mizuno

**Author notes:** Corresponding Author: Tomoyuki Mizuno, PhD, Division of Translational and Clinical Pharmacology, Cincinnati Children’s Hospital Medical Center, Cincinnati, Ohio, USA.

## Abstract

Population pharmacokinetic (PK) model-based Bayesian estimation is widely used for dose individualization, particularly when sample availability is limited. However, its predictive accuracy can be compromised by factors such as misspecified prior information, intra-patient variability, and uncertainties in PK variations. In this study, we developed a hybrid approach that combines machine learning (ML) with population PK-based Bayesian methods to improve the prediction of infliximab concentrations in children with Crohn’s disease. We calculated prediction errors between Bayesian-estimated and observed infliximab concentrations from 292 measurements across 93 patients. Incorporating clinical patient features, we explored various ML algorithms, including linear regression, random forest, support vector regression, neural networks, and XGBoost to correct the Bayesian-based prediction errors. The predictive performance of these ML models was assessed using root mean square error (RMSE) and mean prediction error (MPE) with 5-fold cross-validation. For Bayesian estimation alone, the RMSE and MPE were 4.8 µg/mL and -0.67 µg/mL, respectively. Among the ML algorithms, the XGBoost model demonstrated the best performance, achieving an RMSE of 3.78 ± 0.85 µg/mL and an MPE of -0.03 ± 0.69 µg/mL in 5-fold cross-validation. The ML-corrected Bayesian estimation significantly reduced the absolute prediction error compared to Bayesian estimation alone. This hybrid population PK-ML approach provides a promising framework for improving the predictive performance of Bayesian estimation, with the potential for continuous learning from new clinical data to enhance dose individualization.

**Key points:**

- A new hybrid model combining population pharmacokinetic model-based Bayesian estimation and machine learning significantly improved the accuracy of infliximab concentration predictions in young adult and pediatric patients with Crohn’s disease.
- The developed hybrid model can facilitate infliximab individualized dosing by accounting for changes in clinical conditions and patient-specific factors that the conventional Bayesian estimation approach may not address, and can be integrated into precision dosing dashboards, such as RoadMAB, for real-world clinical application.
- This study indicates that model predictive accuracy can be enhanced by combining the Bayesian method with machine learning, even with a relatively small amount of clinical data. This is particularly encouraging for specific populations, such as pediatric patients, where obtaining rich clinical data is challenging.

## 1. INTRODUCTION

Infliximab, a chimeric monoclonal antibody that targets tumor necrosis factor (TNF)-alpha, is crucial for inducing and maintaining both clinical and endoscopic remission in pediatric Crohn’s disease [1]. It is widely utilized as a first-line therapy for moderate to severe pediatric Crohn’s disease and has demonstrated significant efficacy in improving disease outcomes [2]. However, substantial inter-individual variability in pharmacokinetics and treatment response, influenced by biomarkers such as albumin levels, inflammatory burden, and immunogenicity, highlights the need for individualized dosing strategies [3, 4]. Therapeutic drug monitoring/management (TDM) is commonly used to optimize infliximab dosing, ensuring that patients achieve target trough concentrations and experience optimal therapeutic benefit [5, 6]. Proactive TDM, which involves routinely measuring infliximab concentrations to guide dose adjustments, has been shown to improve disease remission rates and enhance the durability of anti-TNF biologics in children [7, 8].

To support individualized dosing, our research group has developed a population pharmacokinetic (PK) model specifically for pediatric patients with Crohn’s disease [9]. This model has been implemented into a precision dosing PK dashboard, RoadMAB, embedded into our electronic health record (EHR) systems, enabling Bayesian estimation-based dose adjustments by clinicians at the bedside [10]. While Bayesian estimation provides predictions of infliximab concentrations to inform individualized dosing, its predictive performance for future concentrations can be compromised. Factors such as intra-patient variability, changes in disease activity, and alterations in pharmacokinetics over time may limit its accuracy. This discrepancy often stems from time-varying factors not captured by the population PK model, such as changes in target TNF-alpha levels, inflammation, anti-drug antibody development, and growth-related changes in children [11, 12]. These dynamic elements can result in deviations between predicted and actual concentrations, potentially leading to suboptimal dosing and treatment failure [13].

Artificial intelligence (AI) and machine learning (ML) have recently gained attention as promising tools for enhancing TDM and model-informed precision dosing (MIPD) [14, 15]. By identifying complex, nonlinear relationships in clinical data, ML algorithms can correct prediction errors and adapt models to individual patient characteristics. A recent systematic review underscored ML’s promise across various therapeutic areas [16]. Additionally, several investigations have demonstrated the effectiveness of hybrid methods that merge ML and pharmacometrics approaches [15]. For instance, a hybrid approach using ML-based selection of Bayesian prior information reduced the prediction errors in vancomycin concentration [17], while another hybrid approach that integrates PK/PD model parameters into ML model improves predictive performance for chemotherapy-induced neutropenia [18]. Furthermore, the population PK-ML model, which incorporates individual PK parameters as ML features, enhances AUC prediction for vancomycin in patients with sepsis [19]. These findings illustrate how AI and ML can bolster PK predictions and optimize dosing strategies. Nevertheless, despite their demonstrated utility in other therapeutic areas, hybrid models for infliximab precision dosing remain unreported.

In this study, we address this gap by developing a hybrid model that combines Bayesian estimation with ML algorithms to enhance infliximab concentration predictions in pediatric and young adult patients with Crohn’s disease. Our approach utilizes Bayesian estimation for initial predictions, informed by population PK model and observed concentrations, and subsequently integrates ML models to update these predictions based on clinical factors. This hybrid framework aims to correct systematic prediction errors, improve predictive accuracy, and support individualized, adaptive dosing strategies.

## 2. METHODS

### 2.1 Study Design and Data

This study aimed to develop a hybrid approach that combines ML with population PK-based Bayesian methods to improve the predictive accuracy of infliximab concentrations in children with Crohn’s disease. Data was extracted from two clinical studies involving patients aged 2 to 22 years who were receiving infliximab for Crohn’s disease. The first study, REFINE, was a multicenter, prospective, observational study conducted at Cincinnati Children’s Hospital Medical Center (CCHMC), Connecticut Children’s Medical Center, Medical College of Wisconsin, and Nationwide Children’s Hospital between 2014 and 2019. The second study, APPDASH, enrolled patients at Cincinnati Children’s Hospital Medical Center between 2019 and 2021. Both studies were approved by the respective Institutional Review Boards. Patients with two consecutive trough infliximab concentrations exceeding 0.4 µg/mL (lower limit of quantification) were included, yielding 292 pairs of successive concentrations from 93 children. The first concentration (C_Baye_) for each patient was used for Bayesian estimation, while the second concentration (C_obs_) was used to assess prediction performance. The demographic and dosing data are summarized in Table 1.

**Table 1.** Demographic and dosing data in real-world clinical patients.

|  |  |
| --- | --- |
| Number of patients, n | 93 |
| Age, years (median, range) | 13.1 (1.4 - 20.2) |
| Female, n | 46 (49.5%) |
| Weight, kg (median, range) | 38.6 (9.1 - 91.4) |
| Number of successive infliximab trough concentrations, n | 292 |
| Number of infusions at the time of C<sub>Baye</sub> collection (median, range) | 7 (5 - 18) |
| Dose at the time of C<sub>Baye</sub> collection, mg (median, range) | 300 (100 - 1000) |
| Dose at the time of C<sub>obs</sub> collection, mg (median, range) | 400 (100 - 1000) |
| Dose intervals, weeks (median, range) | 6 (2 - 17) |
| Time between the C<sub>Baye</sub> and the C<sub>obs</sub> , weeks (median, range) | 6 (2 - 16) |

C<sub>Baye</sub>, the concentration that was used for Bayesian estimation; C<sub>obs</sub>, the concentration to be predicted (i.e. one that was used to assess prediction performance).

### 2.2 Bayesian Estimation with Population Pharmacokinetic Model

Bayesian estimation was performed using a previously developed population PK model for pediatric patients [9]. The mapbayr R package [20] was used to implement maximum a posteriori Bayesian estimation, predicting individual PK parameter estimates. All dosing history and covariates including body weight, serum albumin (ALB), erythrocyte sedimentation rate (ESR), neutrophil CD64 (nCD64), and antibody to infliximab (ATI) were used for Bayesian estimation. Missing covariate data was imputed by reference values of the population PK model. Prediction errors were calculated as the difference between the Bayesian-estimated trough concentration (C_pred_) and the observed trough concentration (C_obs_) at the next dose as follows:

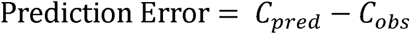

### 2.3 Development of Machine Learning Models

ML models were built to predict prediction errors derived from the Bayesian estimation. Several ML algorithms were explored, including linear regression, random forest, support vector regression, neural networks, and XGBoost. The training data included patient-specific covariates, demographic information, and Bayesian-estimated PK parameters. Pearson correlation coefficients were calculated to identify patient features associated with the prediction errors. Hyperparameter tuning was performed with a grid search using five-fold cross-validation to optimize model performance.

### 2.4 Hybrid Model Integration

The hybrid model integrated Bayesian PK predictions with ML model-based corrections to improve infliximab concentration prediction, as depicted in Figure 1. The ML models, trained on the prediction errors, aimed to capture systematic biases and unexplained variability.

**Figure 1.**
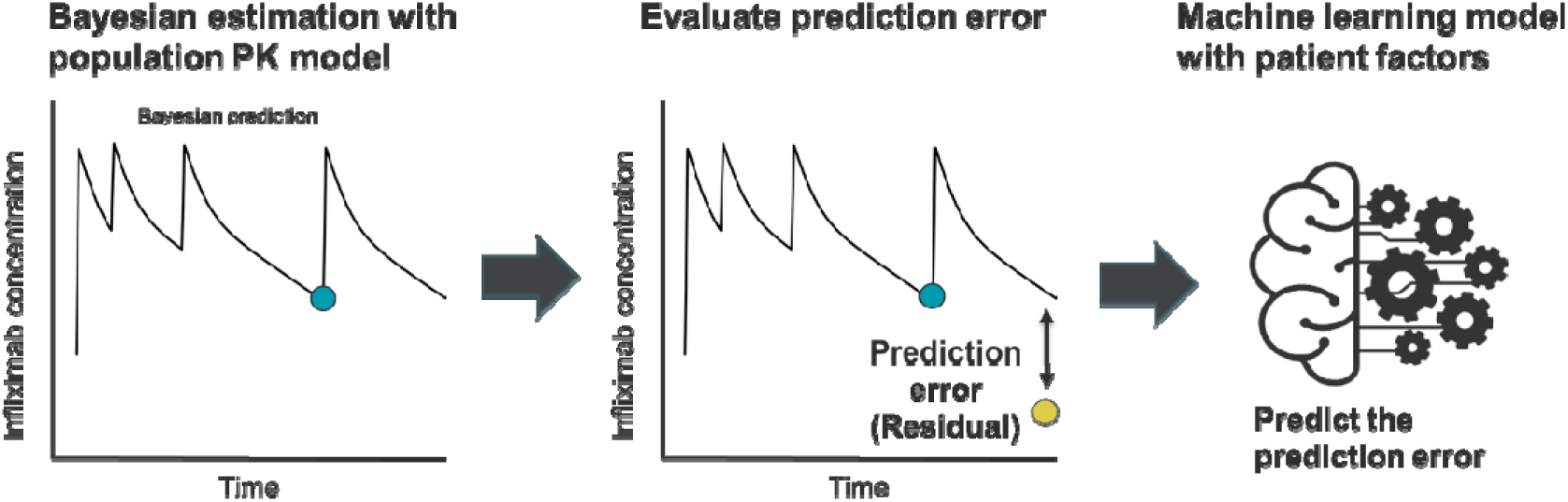
Concept of hybrid population PK-machine learning modeling approach. The blue dot indicates the concentration used for Bayesian estimation (C_Baye_) and the yellow dot indicates concentration used to assess prediction performance (C_obs_).

Ensemble predictions were generated by adding the ML-based predicted error to the Bayesian PK prediction as follows:

where C_pred_ is the predicted infliximab concentration by Bayesian estimation. Prediction error_M_ is a prediction error predicted using the ML models.

### 2.5 Evaluation of Prediction Performance

A five-fold cross-validation approach was employed to robustly evaluate model performance. Prediction accuracy was assessed using two metrics: mean prediction error (MPE) and root mean squared error (RMSE). The equations are defined as follows, where C_pred_ is the predicted trough concentration and C_obs_ is the observed trough concentration:

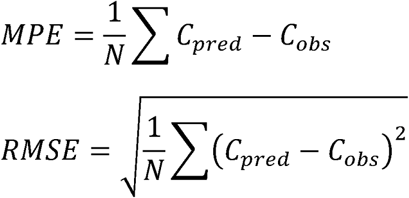

To statistically evaluate the prediction performance between the hybrid model and the Bayesian method, absolute prediction error (APE, |C_pred D_ C_obs_|) generated by 5-fold cross-validation were compared with those estimated by the Bayesian method using the Wilcoxon signed-rank test. A p-value less than 0.05 was considered statistically significant.

### 2.6 Software

All analyses were performed using RStudio (version 2024.04.0) with R (version 4.4.0) and Rtools 4.4. XGBoost models were developed using the xgboost package (version 1.7.8.1), while PK simulations were conducted with the mrgsolve package (version 1.5.2). Bayesian estimation based on the population PK model was carried out using the mapbayr package (version 0.10.0). For support vector regression, the e1071 package (version 1.7-16) was used; neural networks were implemented with the nnet package (version 7.3-19); and random forest models were built using the randomForest package (version 4.7-1.2).

## 3. RESULTS

### 3.1 Predictive Performance of Bayesian Estimation

For Bayesian estimation alone, the RMSE was 4.8 µg/mL, and the MPE was -0.67 µg/mL, indicating room for improvement in predictive performance. Prediction errors showed significant associations with various patient-specific factors (Table 2), including the observed infliximab concentration used for Bayesian estimation (prior trough; r = -0.47, p < 0.001), deviation from typical clearance (ETA for CL; r = -0.14, p = 0.0168), time between the C_Baye_ and the C_obs_ (r=0.192, p<0.001), ESR (r = -0.15, p = 0.015), anti-infliximab antibodies (r = -0.19, p = 0.001), weight change (r = -0.16, p = 0.005), and changes in ESR during the previous dosing interval (r = 0.23, p < 0.001). These findings suggest that incorporating these factors into the model could help improve the predictive performance of Bayesian estimation. Accordingly, the seven factors with *p* < 0.05 were selected as input features for the ML models.

**Table 2.** Patient-specific factors associated with prediction errors of Bayesian estimation alone.

| **Feature** | **r** | **P-value** |
| --- | --- | --- |
| **Observed infliximab concentration ($C_{\text{Baye}}$)**<br>**for Bayesian estimation** | -0.47 | < 0.001 |
| **Predicted infliximab concentration ($C_{\text{pred}}$)**<br>**by Bayesian estimation** | 0.1 | 0.087 |
| **Estimated CL (L/h)** | -0.07 | 0.218 |
| **Estimated CL (L/h/70kg)** | -0.114 | 0.052 |
| **ETA for CL**<br>**(Deviation to mean CL)** | -0.14 | 0.017 |
| **Estimated V1 (L)** | 0.092 | 0.12 |
| **Cumulative dose** | 0.05 | 0.425 |
| **Cumulative dose per time** | 0.07 | 0.254 |
| **Cumulative dose per kg per time** | -0.05 | 0.439 |
| **Dose at the time of $C_{\text{Baye}}$** | 0.08 | 0.15 |
| **Dose at the time of $C_{\text{obs}}$** | 0.05 | 0.426 |
| **Dose at the time of $C_{\text{Baye}}$ per kg** | -0.01 | 0.812 |
| **Dose at the time of $C_{\text{obs}}$ per kg** | -0.05 | 0.366 |
| **Time at the $C_{\text{Baye}}$** | 0.01 | 0.814 |
| **Time at the $C_{\text{obs}}$** | 0.02 | 0.763 |
| **Time between the $C_{\text{Baye}}$ and the $C_{\text{obs}}$** | 0.192 | < 0.001 |
| **WT at the time of $C_{\text{Baye}}$** | 0.1 | 0.085 |
| **WT at the time of $C_{\text{obs}}$** | 0.09 | 0.132 |
| **ALB at the time of $C_{\text{Baye}}$** | 0.03 | 0.64 |
| **ALB at the time of $C_{\text{obs}}$** | -0.01 | 0.904 |
| **ESR at the time of $C_{\text{Baye}}$** | -0.14 | 0.015 |
| **ESR at the time of $C_{\text{obs}}$** | 0.04 | 0.464 |
| **nCD64 at the time of $C_{\text{Baye}}$** | -0.01 | 0.834 |
| **nCD64 at at the time of $C_{\text{obs}}$** | 0.07 | 0.25 |
| **ATI at the time of $C_{\text{Baye}}$** | -0.19 | 0.001 |
| **ΔALB** | -0.04 | 0.463 |
| **ΔESR** | 0.23 | < 0.001 |
| **ΔnCD64** | 0.08 | 0.173 |
| ΔWT | -0.16 | 0.005 |
| ΔAMT | -0.1 | 0.088 |
| ΔTIME | 0.04 | 0.528 |

$C_{\text{Baye}}$ , the concentration that was used for Bayesian estimation; $C_{\text{pred}}$ , Bayesian-estimated concentration; $C_{\text{obs}}$ , the concentration to be predicted (i.e. one that was used to assess prediction performance); $r$ : Pearson correlation coefficient, ATI: anti-infliximab antibody, ALB: serum albumin, ESR: erythrocyte sedimentation rate, ETA for CL: deviation from mean CL, nCD64: neutrophil CD64, WT: body weight, and $\Delta$ : Difference between $C_{\text{Baye}}$ and $C_{\text{obs}}$ .

### 3.2 Evaluation of Predictive Performance of the Hybrid Model

Among the ML algorithms evaluated with selected features, the XGBoost model showed the lowest RMSE and MPE in five-fold cross-validation (Figure 2). The XGBoost model-corrected Bayesian estimation achieved a RMSE (mean ± standard deviation) of 3.78 ± 0.85 µg/mL and an MPE (mean ± standard deviation) of -0.03 ± 0.69 µg/mL, representing an improvement over Bayesian estimation alone. The reduced RMSE and near-zero MPE highlight both enhanced accuracy and minimal bias. The observed and predicted concentration plots for Bayesian estimation alone and hybrid models are shown in Figure 3. Additionally, the Wilcoxon signed-rank test revealed a significant reduction in absolute prediction errors (APE) with the XGBoost-corrected Bayesian estimation compared to the Bayesian estimation alone (Median APE: 1.07 and 1.15 µg/mL, p = 0.013).

**Figure 2.**
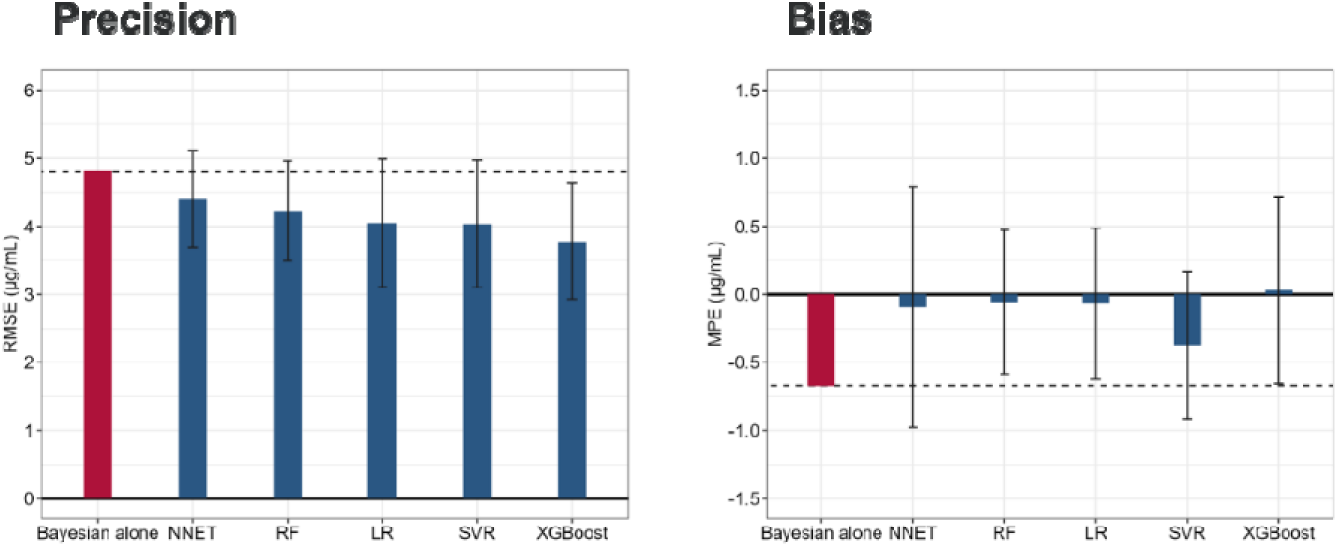
Comparison of predictive performance between Bayesian estimation alone and hybrid models. RMSE, root mean squared error; MPA, mean prediction error; NNET, neural networks; RF, random forest; LR, linear regression; SVR, support vector regression.

**Figure 3.**
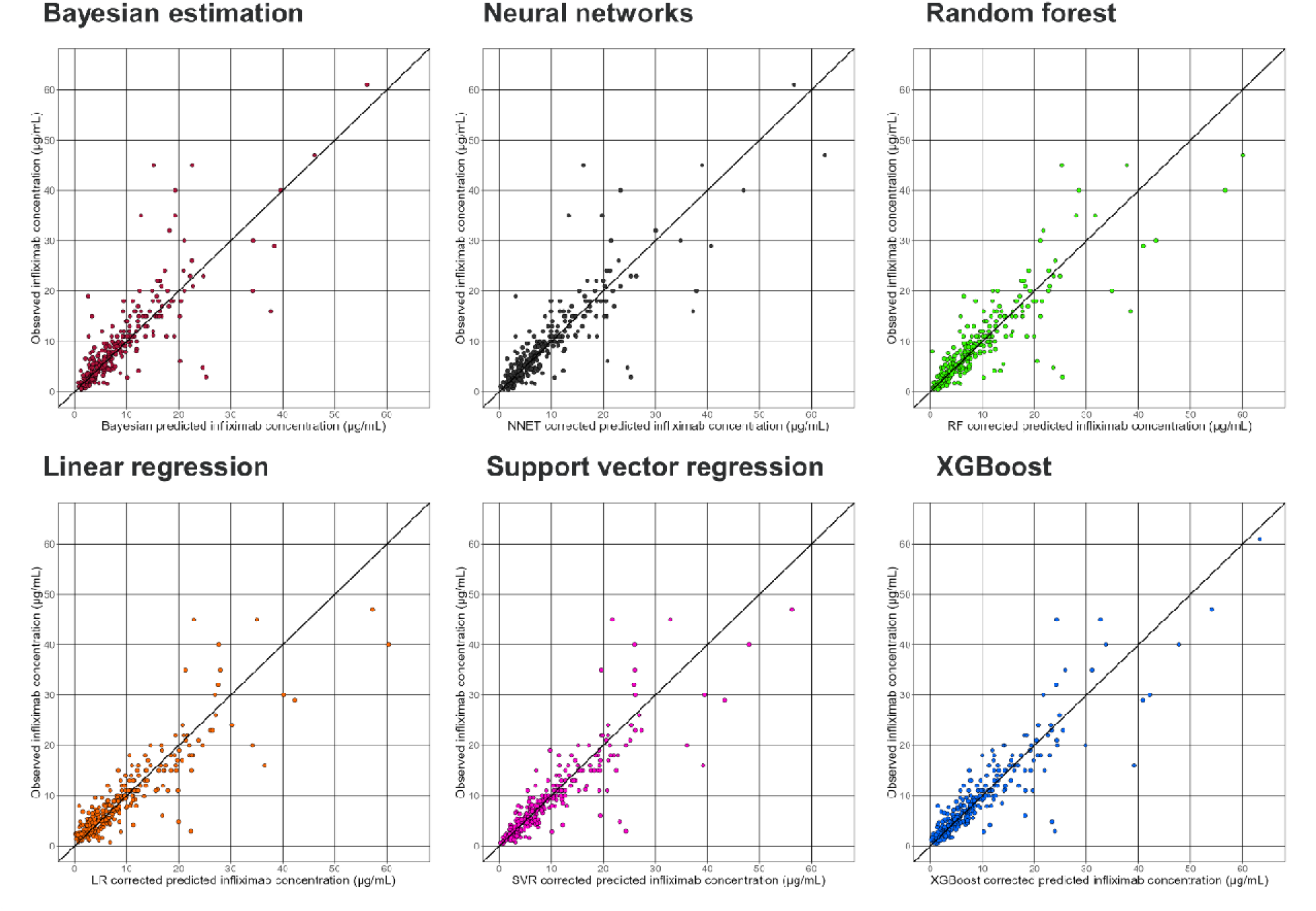
Observation versus prediction plots for Bayesian estimation alone and hybrid models.

### 3.3 Representative Cases

To illustrate the effectiveness of the hybrid model, representative cases from the real-world dataset are presented in Figure 4. In one example (Figure 4, Case 1), the Bayesian-predicted profile (black solid line) overestimated the observed concentration at the subsequent dose (yellow dot), whereas the XGBoost-corrected prediction (pink dot) more closely matched the actual value. In another example (Figure 4, Case 2), the Bayesian method underestimated the observed concentration, and again, the XGBoost correction improved prediction accuracy. Collectively, these examples underscore the hybrid model’s potential to enhance infliximab concentration estimation. Furthermore, as the hybrid model prediction is not time-concentration profiles, PK profiles were then refined using Bayesian estimation under the assumption of inter-occasional variability (20%), as shown in Figure 5. Comparing the initial predicted profiles with Bayesian estimation (Black lines in Figure 5), the refined PK profiles based on the hybrid model predictions (Blue lines) aligned more closely with the observed concentrations. These refined profiles can be used to optimize dosing regimens, enabling more precise and individualized patient care.

**Figure 4.**
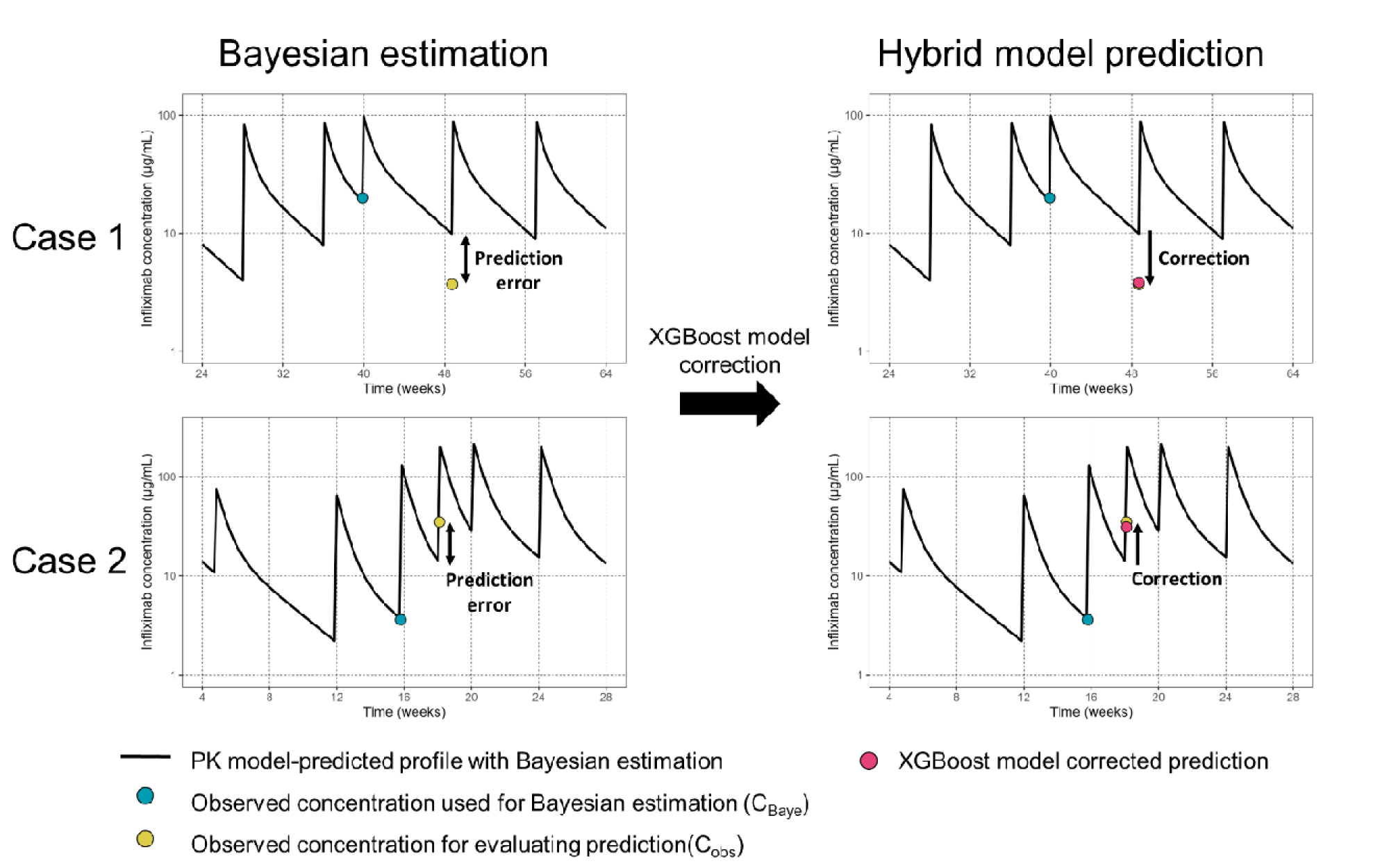
Bayesian predicted PK profiles and Hybrid model-based predictions in representative real-world two cases.

**Figure 5.**
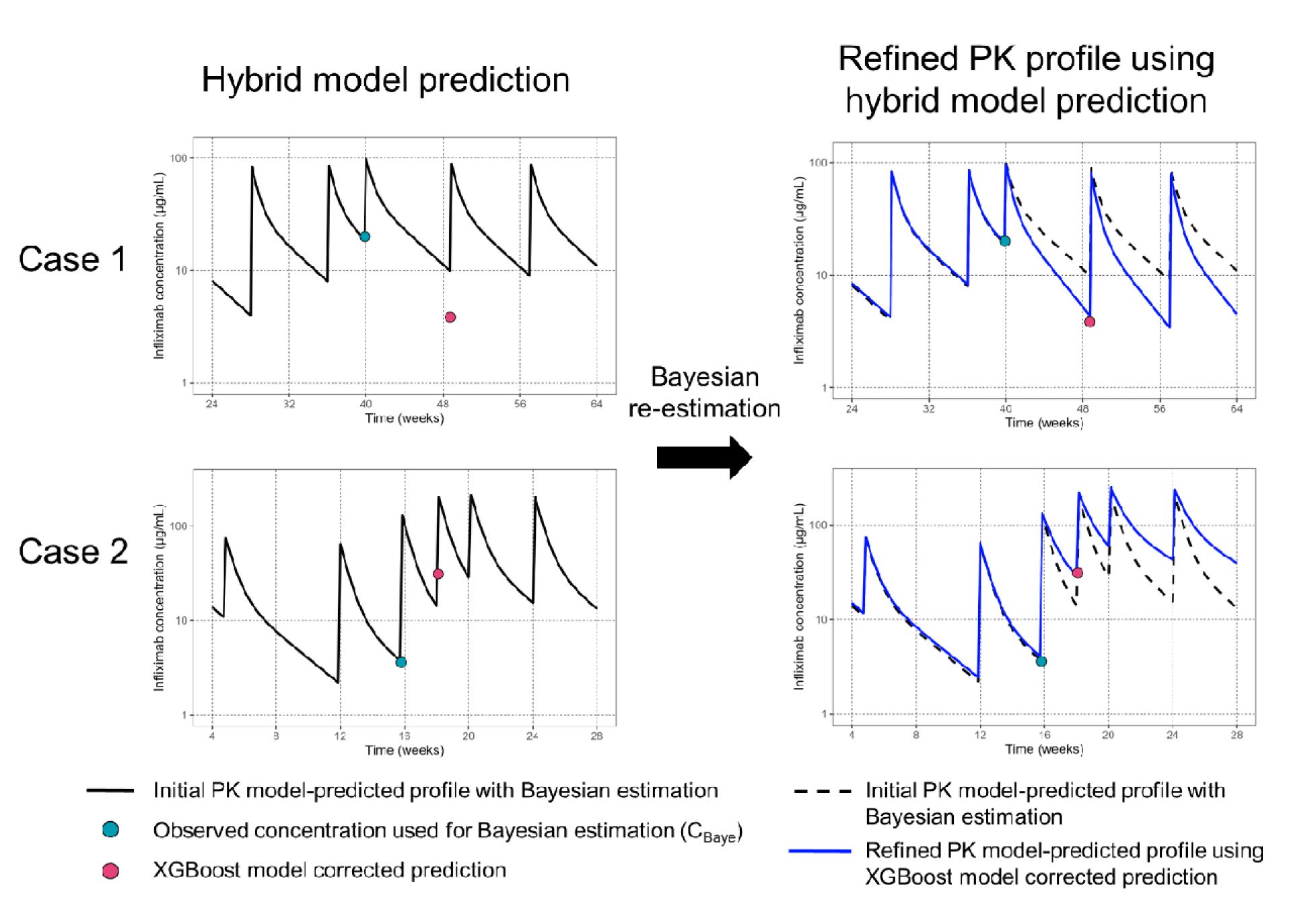
Refinement of model predictions with Bayesian estimation using hybrid model-based predictions in representative real-world two cases.

## 4. DISCUSSION

This study introduced the hybrid model combining population PK-based Bayesian estimation with ML to predict infliximab concentrations in pediatric Crohn’s disease. While Bayesian estimation is widely used as a valuable tool for dose individualization [21], the predicted performance sometimes needs to be improved, especially when patient’s conditions change over time. This study demonstrated that the ML-based complementation for model prediction, especially through XGBoost, significantly enhanced overall predictive performance with relatively small, sparse data. These findings highlight the potential of integrating Bayesian estimation with ML to provide more precise and individualized dosing recommendations for pediatric infliximab therapy.

Although Bayesian estimation provides reasonable predictions of infliximab concentration to inform optimal dosing in children, it still exhibits substantial prediction errors influenced by patient-specific factors [22]. In this study, we identified several factors associated with discrepant predictions, such as prior infliximab concentrations (C_Baye_), clearance variations, time between the C_Baye_ and the C_obs_, ESR, anti-infliximab antibodies, and changes in weight and inflammation. These results suggest that traditional population PK models do not fully account for variability caused by dynamic disease factors (e.g., TNF-alpha fluctuations, inflammation) or pediatric growth. By incorporating ML corrections, the hybrid model successfully addressed these gaps, significantly improving predictive performance and emphasizing the value of integrating additional clinical factors into Bayesian estimation.

Among the ML algorithms evaluated, XGBoost demonstrated the strongest performance. Its ability to model complex, non-linear relationships—often missed by traditional PK models—makes it well-suited for precision dosing tasks. In previous studies, XGBoost has been reported to be effectively applied to scenarios for precision dosing. Tootooni et al used XGBoost model for optimizing vancomycin dosing in critically ill patients by identifying key factors like kidney function and prior drug levels [23]. Similarly, it has been used in methotrexate therapy for pediatric oncology to predict drug clearance and adjust dosing regimens based on liver function and enzyme activity, further establishing its relevance in precision dosing research [24].

A major challenge in developing robust ML models is the need for large datasets to account for variability and ensure generalizability [25]. In specific populations such as pediatrics, gathering such extensive data is often hindered by logistical and clinical constraints [26]. The hybrid approach presented here addresses these limitations by correcting prediction errors based on observed concentrations after initial Bayesian model development. This method leverages knowledge from population PK modeling and available clinical data to enhance performance without requiring vast datasets, making it adaptable across various model-informed precision dosing strategies.

Another significant strength of the hybrid approach is its capacity for continuous learning [27]. By integrating with a precision dosing dashboard [10], the model can update its predictions as new patient data becomes available, thereby refining dosing recommendations in real time. This dynamic adaptability is especially critical in pediatrics, where rapid growth and disease progression necessitate frequent dosing adjustments. By incorporating real-time clinical data, the hybrid model not only refines PK profiles but also identifies new factors influencing pharmacokinetics, enabling more personalized therapy.

Despite these promising advancements, the study has limitations. The relatively small sample size and reliance on data from only two clinical cohorts may limit its generalizability. Additionally, the model has yet to undergo external validation, a crucial step for confirming its applicability and robustness across diverse patient populations and clinical settings. Future research should focus on external validation and the incorporation of larger, more diverse datasets. Notably, the developed hybrid ML model can be integrated into our clinical dashboard, RoadMAB [10], which is designed to implement model-informed precision dosing at the bedside. With this integration, the dashboard could make complex pharmacokinetic predictions more accessible to clinicians and ultimately facilitate improved patient care.

## 5. CONCLUSION

The hybrid model combining Bayesian estimation with ML offers a novel and flexible strategy for precision dosing of infliximab in pediatric Crohn’s disease. By addressing prediction errors and individual variability, this approach facilitates personalized dosing for pediatric patients with Crohn’s disease and other conditions requiring individualized dosing strategies.

## Funding

This work was supported by the National Institute of Diabetes and Digestive and Kidney Diseases at the National Institutes of Health [grant number DK132408], Crohn’s and Colitis Foundation, and the Cincinnati Children’s Research Foundation.

## Conflicts of Interest

J. S. H. is on advisory boards for Janssen, Abbvie, Lilly, and Genentech and is a consultant for Takeda and Pfizer. The other authors report no conflicts of interest.

## Data availability statements

The datasets analyzed in this study are accessible upon reasonable request by contacting the corresponding author.

## Code availability

The model codes are accessible upon reasonable request by contacting the corresponding author.

## Ethics approval

This study was conducted in accordance with the Declaration of Helsinki. The REFINE study received approval from multiple Institutional Review Boards (IRBs), including Cincinnati Children’s Hospital Medical Center (CCHMC), Nationwide Children’s Hospital, Medical College of Wisconsin, and Connecticut Children’s Medical Center. Additionally, the CCHMC IRB approved the APPDASH study.

## Consent to Participate

Written informed consent was obtained from each participant, or from a parent or legal guardian, as appropriate. A waiver of consent was granted when applicable.

## Consent for publication

Not applicable

## Author Contributions

K.I., P.M., and T.M. wrote the manuscript; K.I., P.M., and T.M. designed the research; K.I., P.M., J.R., BMB, JDN, JSH and T.M. performed the research; and K.I. and T.M. analyzed the data.

